# The ability of the LIMD1 and TRIP6 LIM domains to bind to f-actin under strain is critical for their tension dependent localization to adherens junctions and association with the Hippo pathway kinase LATS1

**DOI:** 10.1101/2023.10.12.562097

**Authors:** Samriddha Ray, Chamika DeSilva, Ishani Dasgupta, Sebastian Mana-Capelli, Natasha Cruz-Calderon, Dannel McCollum

## Abstract

A key step in regulation of Hippo pathway signaling in response to mechanical tension is recruitment of the LIM domain proteins TRIP6 and LIMD1 to adherens junctions. Mechanical tension also triggers TRIP6 and LIMD1 to bind and inhibit the Hippo pathway kinase LATS1. How TRIP6 and LIMD1 are recruited to adherens junctions in response to tension is not clear, but previous studies suggested that they could be regulated by the known mechanosensory proteins *α*-catenin and vinculin at adherens junctions. We found that the three LIM domains of TRIP6 and LIMD1 are necessary and sufficient for tension-dependent localization to adherens junctions. The LIM domains of TRIP6, LIMD1, and certain other LIM domain proteins have been shown to bind to actin networks under strain/tension. Consistent with this, we show that TRIP6 and LIMD1 colocalize with the ends of actin fibers at adherens junctions. Point mutations in a key conserved residue in each LIM domain that are predicted to impair binding to f-actin under strain inhibits TRIP6 and LIMD1 localization to adherens junctions and their ability to bind to and recruit LATS1 to adherens junctions. Together these results show that the ability of TRIP6 and LIMD1 to bind to strained actin underlies their ability to localize to adherens junctions and regulate LATS1 in response to mechanical tension.

## Introduction

The ability of cells to sense and respond to mechanical cues is essential to preserve cell integrity and make cell fate decisions regarding proliferation, survival, and differentiation (1, 2). Cells deal with a diverse array of mechanical forces. These forces can be external such as shear stress, stretch, or compression, forces from neighboring cells, or internal forces from their own contractile apparatus. Other factors such as substrate stiffness and cell crowding also have profound effects on cell behavior. For example, epithelial cells at low density are subject to greater pulling forces from neighboring cells than cells at high density (3). The ability of normal cells to stop proliferating once they reach a certain density is mediated at least in part by regulation of the Hippo pathway by pulling forces/tension from neighboring cells that are sensed at adherens junctions (3-5). Adherens junctions are a good place to sense mechanical tension in tissues, since they act as a bridge to connect the cells internal actomyosin contractile apparatus to neighboring cells. The transmembrane cadherin proteins bind to cadherins on neighboring cells, and their cytoplasmic tails bind to various proteins including α-catenin which binds directly and indirectly to the actomyosin cytoskeleton.

At a molecular level, Hippo signaling (6) has been shown to be regulated by tension at adherens junctions through two related LIM domain proteins called TRIP6 (4) and LIMD1 (5). Both TRIP6 and LIMD1 localize to adherens junctions in response to mechanical tension where they recruit and inhibit the Hippo pathway kinases LATS1/2. The details of how both of these proteins are recruited to adherens junctions by mechanical tension are poorly understood. One candidate mechanism is through the known adherens junction mechanosensory protein *α*-catenin (7), which can directly sense mechanical tension across epithelial sheets of cells caused by cell contraction and/or by external forces (8-10). The *α*-catenin protein connects transmembrane cadherin proteins to intracellular actin filaments. Pulling forces from neighboring cells and the internal actomyosin cytoskeleton causes a conformational change in *α*-catenin allowing it to bind and activate vinculin (8, 11). Activated vinculin then binds F-actin and other effector proteins to reinforce the adherens junction (12, 13). Various lines of evidence are consistent with *α*-catenin recruiting LIMD1 and TRIP6 to adherens junctions in response to tension. One study showed that *α*-catenin and LIMD1 are in close proximity of each other at adherens junctions and knockdown of *α*-catenin with siRNA disrupts LIMD1 localization, although it also disrupts adherens junctions (5). A different study showed that TRIP6 binds to vinculin and its localization to adherens junctions depends on vinculin and mechanical tension (4). These results could be explained by a model where in response to mechanical tension *α*-catenin is activated and binds to and activates vinculin. Then TRIP6 and LIMD1 are recruited by *α*-catenin, vinculin, or other downstream effectors. Although this model is consistent with previous data, it is still not clear whether *α*-catenin and vinculin directly regulate TRIP6 and LIMD1 localization and activation, or if they are recruited through other tension dependent mechanisms. Here we show that vinculin binding to TRIP6 is not required for recruitment of TRIP6 to adherens junctions in response to mechanical tension. Instead, we show that the LIM domains of both TRIP6 and LIMD1 are sufficient for their mechano-sensitive localization to adherens junctions and their ability to sense tension is required for regulation of LATS1/2.

## Results

### Does *α*-catenin trigger vinculin binding to TRIP6?

To determine whether recruitment of TRIP6 to adherens junctions in response to mechanical strain was directly regulated by *α*-catenin and vinculin, we investigated whether TRIP6-vinculin binding depends on *α*-catenin. We found that TRIP6-vinculin binding was completely lost in *α*-catenin knockout HEK293A cells compared to wild-type cells (Figure 1A). We also observed that cell-cell adhesion was impaired and TRIP6 did not localize to cell-cell junctions in *α*-catenin knockout cells, suggesting that the lack of interaction between TRIP6 and vinculin may reflect a general perturbation of adherens junctions in these cells (Figure 1B). To test whether *α*-catenin promoted vinculin-TRIP6 binding directly, we examined whether the vinculin-T12 mutant (14), which bypasses the requirement for *α*-catenin for activation, can rescue the vinculin-TRIP6 binding defect in *α*-catenin null cells. These experiments showed that vinculin-T12 only very weakly rescued the ability to bind TRIP6 in *α*-catenin knockout cells (Figure 1A). These results show that although *α*-catenin is required for vinculin-TRIP6 binding, the requirement may be indirect, and depend mainly on other functions of *α*-catenin such as junction assembly, actin recruitment, and/or ability to generate tension at junctions (15).

**Figure 1.**
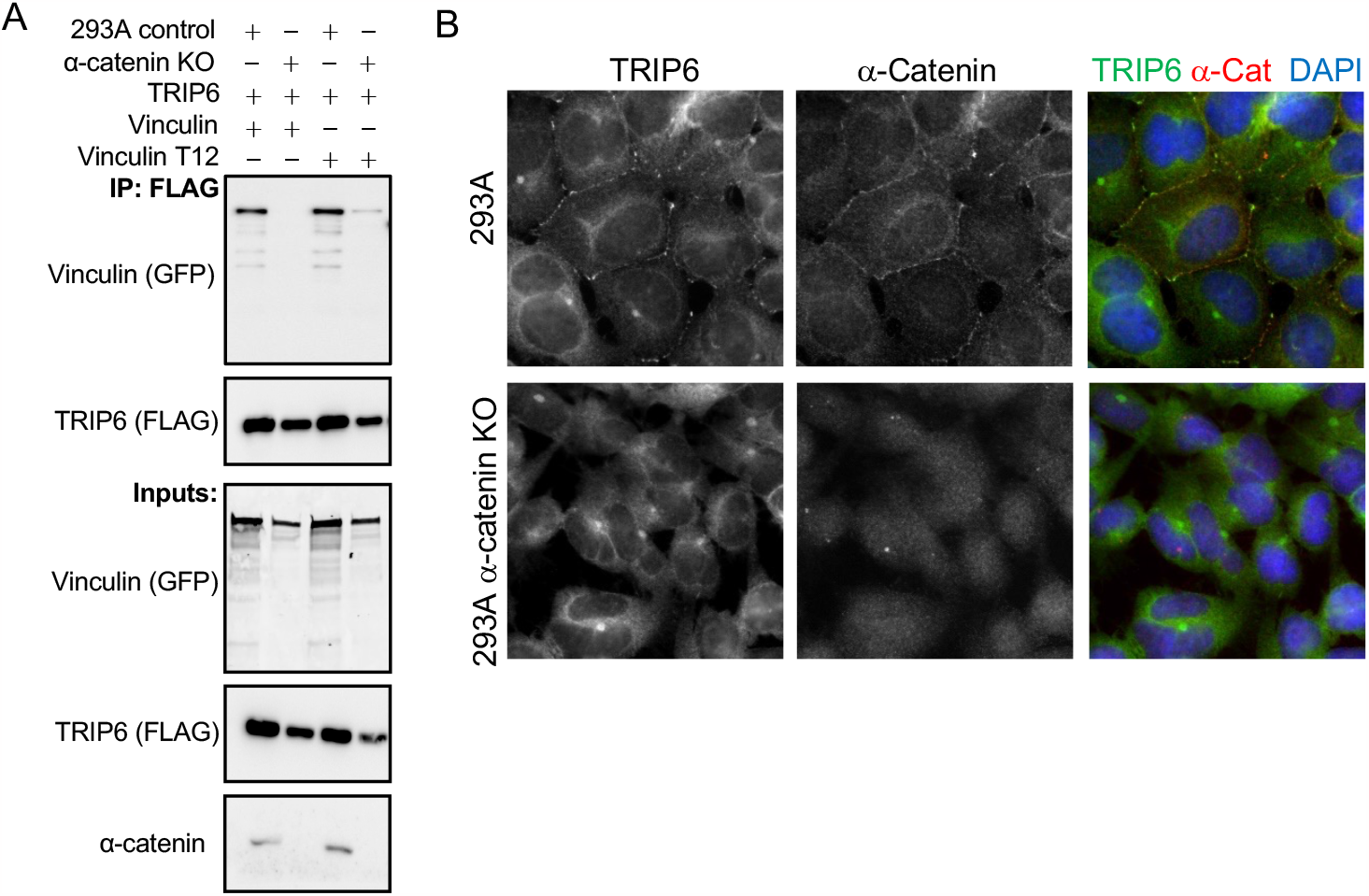
The requirement for *α*-E catenin for TRIP6-Vinculin binding and TRIP6 adherens junction localization may be indirect. (A) HEK293A (293A control) or HEK293A cells with *α*-E catenin inactivated using CRISPR (*α*-catenin KO) were transfected with a FLAG-TRIP6 expressing plasmid (TRIP6) and either a plasmid for expressing GFP-vinculin (Vinculin) or the GFP-vinculin-T12 mutant (Vinculin T12). FLAG-TRIP6 immune complexes were isolated from cell lysates and analyzed by western blotting using antibodies against GFP (Vinculin (GFP)) and FLAG (TRIP6 (FLAG)). Cell lysates (Inputs) were analyzed using antibodies against GFP (Vinculin (GFP)), FLAG (TRIP6 (FLAG)), and *α*-E catenin (*α*-catenin). (B) HEK293A (293A) or HEK293A cells with *α*-E catenin inactivated (293A *α*-catenin KO) using CRISPR were stained using anti-TRIP6 and anti-*α*-catenin antibodies. Merged image shows TRIP6 (green), *α*-E catenin (red), and DNA (blue).

### Vinculin binding to TRIP6 is not required for TRIP6-LATS1/2 binding

Our previous study showed that knockdown of vinculin using siRNA reduces binding of TRIP6 to LATS1 and localization of both proteins to adherens junctions, suggesting that vinculin binding to TRIP6 might recruit TRIP6 to adherens junctions and enhance its ability to bind LATS1/2. To test this possibility, we first investigated which region(s) of TRIP6 (Figure 2A) are required for binding to each protein. We observed that vinculin bound to the amino-terminal (non-LIM domain) half of TRIP6 (Figure 2C). As observed previously (4), LATS2 bound the carboxy-terminal LIM domain-containing half of TRIP6 (Figure 2D). This binding depended largely on the LIM domains, since deletion of one or more LIM domains significantly reduced binding (Figure 2D). Interestingly, vinculin and LATS2 bind better to the amino (amino acids 1-277) and carboxy terminal (amino acids 278-476) halves of TRIP6 respectively than to full length TRIP6 (Figure 2C-D) suggesting that full length TRIP6 may be in a partially closed conformation. Similar results were observed with the related LIM domain protein Zyxin, where in vitro experiments showed that the 3 LIM domains (but not full length Zyxin) bound to LATS2 (16). To better understand how vinculin bound to TRIP6, we deleted short blocks of conserved amino acids in the amino-terminus of TRIP6. Two of these deletions (Δ1-9, and Δ252-277) negatively affected TRIP6-vinculin binding, with the Δ252-277 construct completely eliminating binding (Figure 2E).

**Figure 2.**
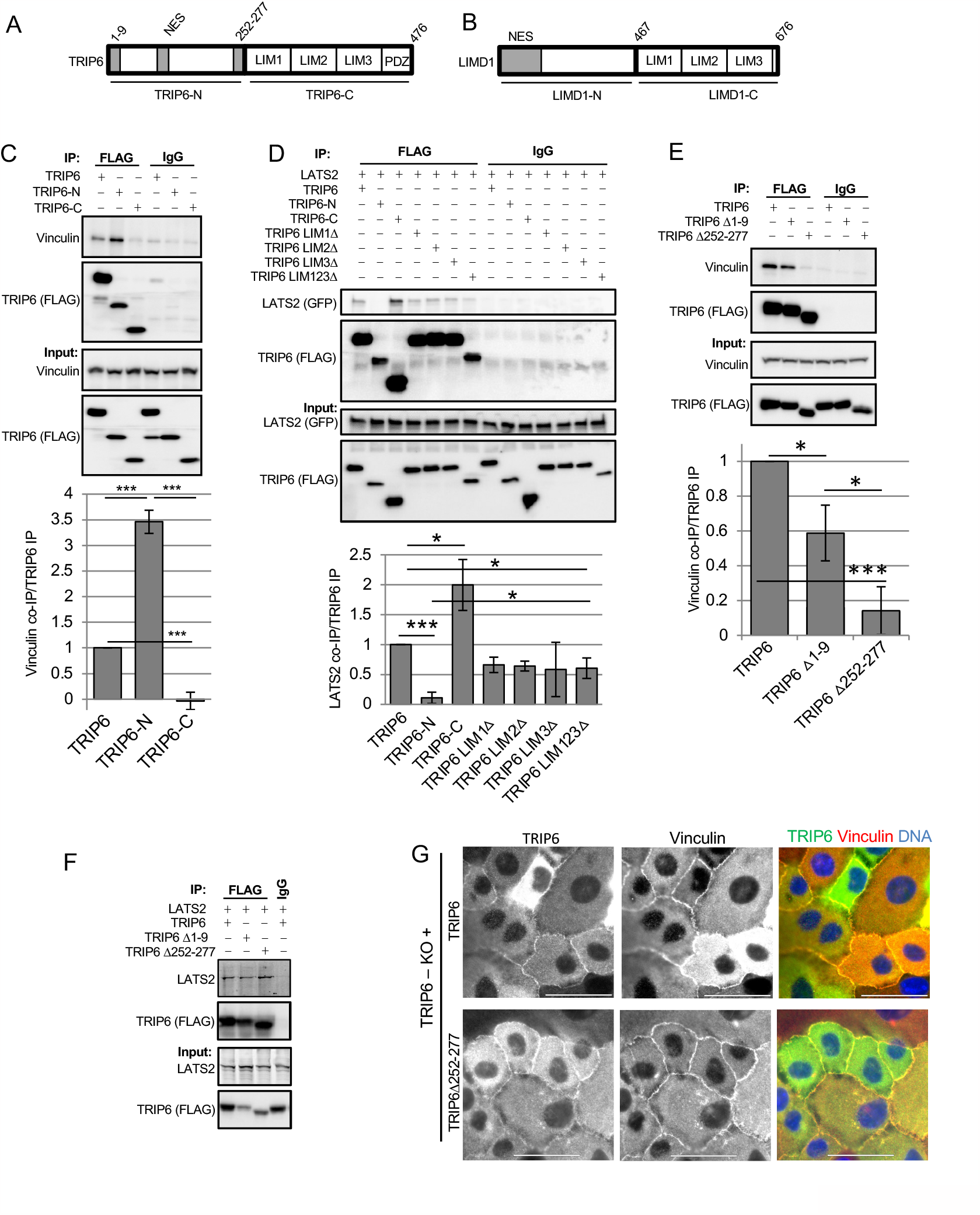
Identification of binding interaction between TRIP6, LATS2, and Vinculin. (A,B) Schematic diagram showing different domains of TRIP6 and LIMD1 (NES: nuclear export signal; LIM: LIM domain; PDZ: PDZ domain binding motif). (C) HEK293 cells were transfected with plasmids expressing vinculin and different FLAG-TRIP6 deletion mutants as indicated. FLAG-TRIP6 immune complexes were isolated from cell lysates and analyzed by western blotting using antibodies against vinculin and FLAG (TRIP6 (FLAG)). Cell lysates (Inputs) were analyzed using antibodies against vinculin and FLAG (TRIP6 (FLAG). Quantification of vinculin levels in TRIP6 immune complexes is shown. (D) HEK293 cells were transfected with plasmids expressing GFP-LATS2 and different FLAG-TRIP6 deletion mutants as indicated. FLAG-TRIP6 immune complexes were isolated from cell lysates and analyzed by western blotting using antibodies against GFP (LATS2 (GFP)) and FLAG (TRIP6 (FLAG)). Cell lysates (Inputs) were analyzed using antibodies against GFP (LATS2 (GFP)) and FLAG (TRIP6 (FLAG)). Quantification of LATS2 levels in TRIP6 immune complexes is shown. (Mean ± SD; n=3; ^*^P≤0.05, ^***^P≤0.001, T-test). (E) HEK293 cells were transfected with plasmids expressing vinculin and different FLAG-TRIP6 deletion mutants as indicated. FLAG-TRIP6 immune complexes were isolated from cell lysates and analyzed by western blotting using antibodies against vinculin and FLAG (TRIP6 (FLAG)). Cell lysates (Inputs) were analyzed using antibodies against vinculin and FLAG (TRIP6 (FLAG)). Quantification of vinculin levels in TRIP6 immune complexes is shown. (Mean ± SD; n=3; ^*^P≤0.05, ^***^P≤0.001, T-test). (F) HEK293 cells were transfected with plasmids expressing GFP-LATS2 and different FLAG-TRIP6 deletion mutants as indicated. FLAG-TRIP6 immune complexes were isolated from cell lysates and analyzed by western blotting using antibodies against GFP (LATS2) and FLAG (TRIP6 (FLAG)). Cell lysates (Inputs) were analyzed using antibodies against GFP (LATS2) and FLAG (TRIP6 (FLAG)). (G) Lentiviral infection was used to stably express GFP fusions of wild-type TRIP6 and TRIP6-Δ252-277 in TRIP6 knockout (TRIP6-KO) MCF10A cells. The indicated cell lines were stained using anti-GFP and anti-vinculin antibodies. Merged image shows TRIP6 (green), vinculin (red), and DNA (blue). Scale bar =50microns.

Interestingly, deletion of amino acids 252-277 did not impair TRIP6 binding to LATS2 (Figure 2F). This is consistent with our previous study (4), which showed that a vinculin mutant with enhanced binding to TRIP6 does not increase binding of TRIP6 to LATS2. Thus, both vinculin mutants that enhance its binding to TRIP6 as well as TRIP6 mutants defective for binding vinculin, do not affect TRIP6 binding to LATS2. Examination of the localization of the TRIP6-Δ252-277 in MCF10A cells where the endogenous copy of TRIP6 had been deleted showed that the TRIP6-Δ252-277 mutant still localizes to adherens junctions (Figure 2G). Together these results show that although vinculin knockdown impairs TRIP6 localization to adherens junctions and binding to LATS1/2 (4), the effect is not through direct binding of vinculin to TRIP6 and may be due to other functions of vinculin such as recruitment of F-actin or other effectors (17).

### The LIM domains of TRIP6 and LIMD1 are critical for tension dependent localization to adherens junctions

To determine which regions of TRIP6 are required for recruitment to adherens junctions, we examined the localization of the amino and carboxy terminal halves of TRIP6 (amino acids 1-277 and 278-476 respectively). Because TRIP6-278-476 lacks the nuclear export signal present in the amino-terminal half of the protein and localizes primarily to the nucleus (not shown) we appended a nuclear export signal to both constructs to assess where they localize in the cytoplasm. TRIP6-1-277 localized to the cytoplasm and did not show any localization to adherens junctions (Figure 3A). In contrast, TRIP6-278-476 localized to adherens junctions (Figure 3A). This localization was still tension dependent since it was disrupted by addition of blebbistatin (Figure 3B). Addition of the vinculin binding site (TRIP6-252-476) did not enhance localization to adherens junctions compared to TRIP6-278-476 (data not shown). The TRIP6-278-476 construct consists of the 3 LIM domains (amino acids 278-472) and a carboxy-terminal PDZ domain binding peptide (amino acids 473-476). To test whether the PDZ domain binding site contributed to adherens junction localization, it was deleted by removing the last 4 amino acids. This construct (TRIP6-278-472) localized to adherens junctions similarly to TRIP6-278-476 (Figure 3A). Thus, the primary targeting information for tension-dependent localization to adherens junctions for TRIP6 is in the 3 LIM domains.

**Figure 3.**
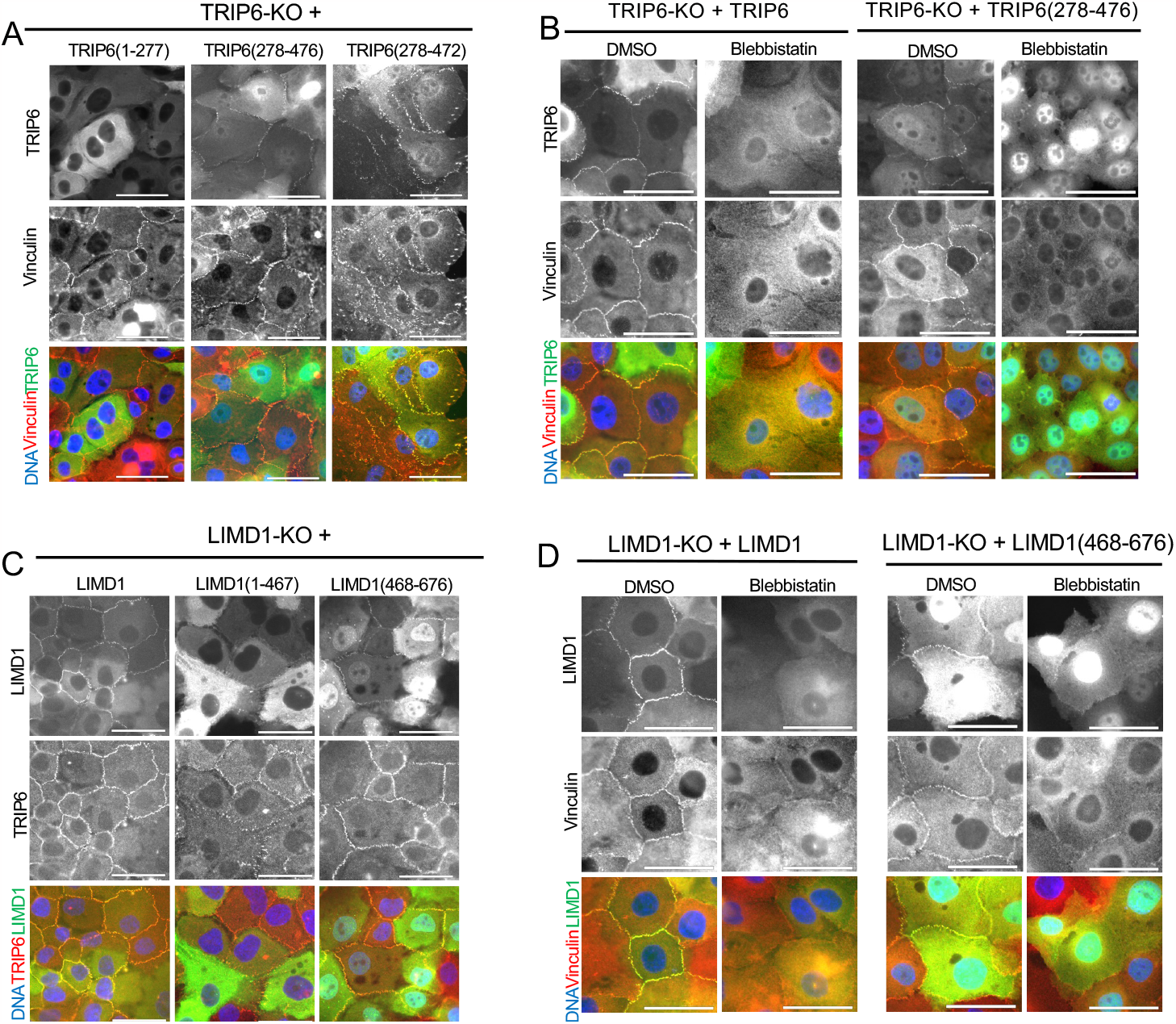
The TRIP6 and LIMD1 LIM domains are necessary and sufficient for tension dependent localization to adherens junctions. Lentiviral infection was used to stably express GFP fusions of wild-type TRIP6, LIMD1 and various deletion mutants of TRIP6 and LIMD1 in TRIP6 knockout (TRIP6-KO) and LIMD1 knockout (LIMD1-KO) MCF10A cells. For (A) and (B), adherens junction localization of indicated TRIP6 constructs was visualized by co-immunostaining using anti-GFP and anti-Vinculin antibody. For (C) and (D), junction localization of indicated LIMD1 constructs was visualized using GFP fluorescence and immunostaining with anti-TRIP6 antibody. Merged images show TRIP6/LIMD1 (green), Vinculin/TRIP6 (red), and DNA (blue). For (B) and (D), indicated cell lines were treated with DMSO (solvent control) or with blebbistatin before fixation and staining. Scale bar =50microns.

Like TRIP6, LIMD1 localizes to adherens junctions and recruits LATS1/2 kinases in response to mechanical tension. To test whether the LIM domains of LIMD1, like those of TRIP6, are sufficient for tension dependent localization of LIMD1 to adherens junctions, we examined the localization of the amino and carboxy terminal halves of LIMD1 (amino acids 1-467 and 468-676 respectively) (Figure 3C) in MCF10A cells where the endogenous copy of LIMD1 had been deleted. Because LIMD1-468-676, which contains the 3 LIM domains of LIMD1, lacks the nuclear export signal present in the amino-terminal half of the protein and localizes primarily to the nucleus (not shown) we appended a nuclear export signal to both constructs to assess where they localize in the cytoplasm. LIMD1-1-467 localized to the cytoplasm, but could occasionally be faintly observed at adherens junctions (Figure 3C). The 3 LIM domains of LIMD1 (LIMD1-468-676) localized to adherens junctions (Figure 3C) almost as well as the full length protein. This localization was still tension dependent since it was disrupted by addition of blebbistatin (Figure 3D). These results show that although the N-terminal region of LIMD1 can weakly localize to adherens junctions, the 3 LIM domains of LIMD1 (as with TRIP6) are the primary driver of tension-dependent localization to adherens junctions.

We next wanted to determine how the LIM domains of TRIP6 and LIMD1 target their respective proteins to adherens junctions. Recent studies have shown that certain classes of LIM domains (including those in TRIP6 and LIMD1) are capable of binding specifically to actin networks under tension/strain (18, 19). Since adherens junctions have strong associations with actin networks, this raised the possibility that the LIM domains of TRIP6 and LIMD1 could be binding and localizing to F-actin under strain at adherens junctions. Indeed, full length TRIP6 and LIMD1, as well as their respective 3 LIM domains alone, colocalize with the ends of actin fibers at adherens junctions (Figure 4A-B). We next tested whether the ability of the LIM domains to sense strained f-actin is critical for TRIP6 and LIMD1 localization. A previous study showed that most of the LIM domains that could sense tension have a conserved phenylalanine in each LIM domain (19). The only exceptions are the 3^rd^LIM domain of zyxin family members (including TRIP6 and LIMD1), which lack the phenylalanine in that position, but have either an adjacent phenylalanine (zyxin) or a tyrosine (TRIP6 and LIMD1). For several proteins (TRIP6 and LIMD1 were not tested) it was shown that the phenylalanine in this position (or the adjacent position for the 3^rd^LIM domain of zyxin) was important for binding to actin under strain. Typically, the phenylalanine had to be mutated in all LIM domains to maximally reduce tension dependent localization of the proteins to F-actin suggesting that each LIM domain contributes additively.

**Figure 4.**
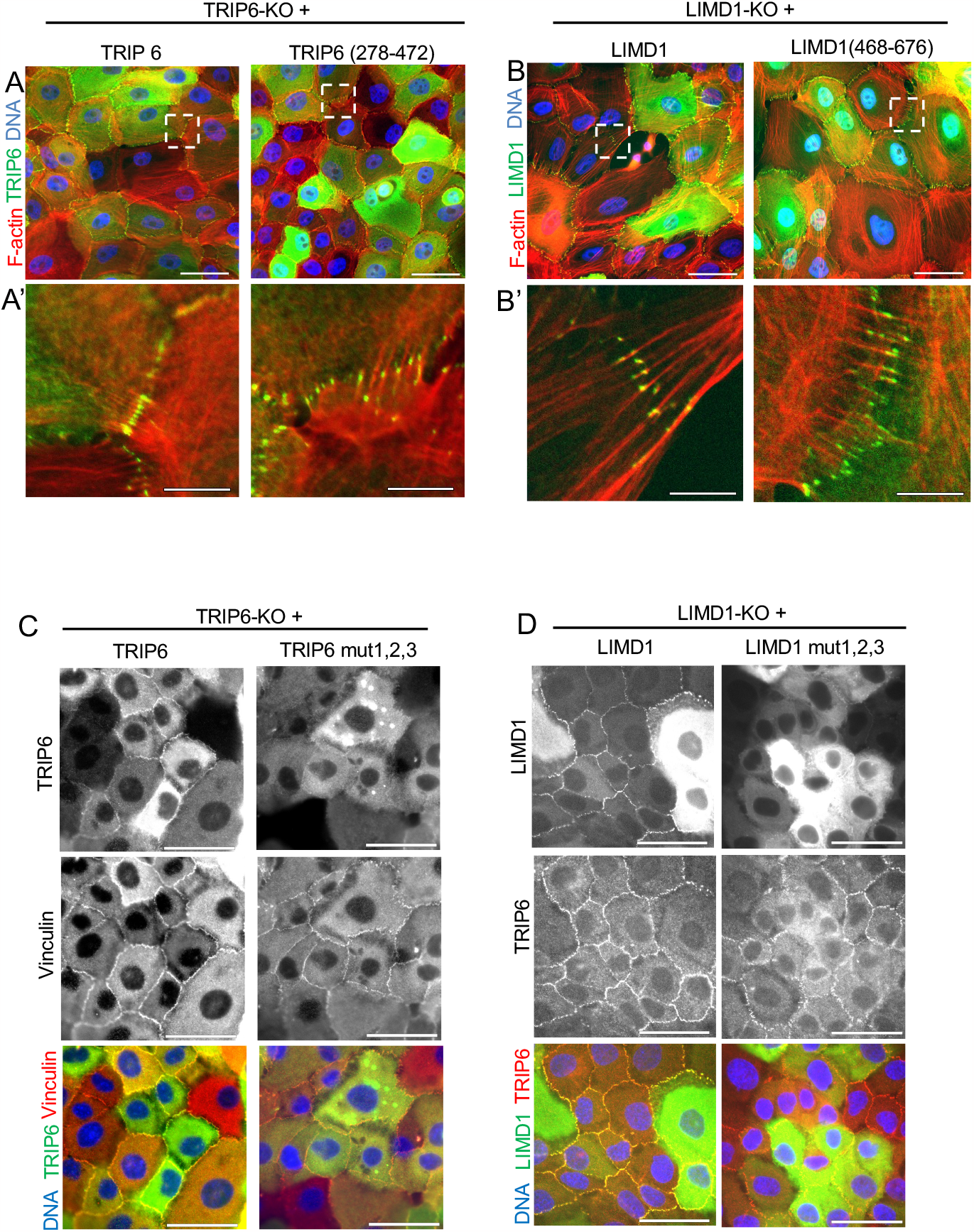
Tension dependent localization of TRIP6 and LIMD1 depends on the ability of their LIM domains to bind f-actin under strain. (A-B) TRIP6 and LIMD1 localize to the ends of actin filaments at adherens junctions. F-actin was visualized by staining with fluorescently tagged phalloidin and localization of GFP constructs was visualized by either using anti-GFP antibodies (for TRIP6 GFP constructs) or using GFP fluorescence (for LIMD1 GFP constructs). Top panels (A,B) scale bars =50 microns. Bottom panels (A’,B’) scale bars = 10 microns, represent blow up views of adherens junctions marked by dashed white box in respective top panel. (C-D) Mutations predicted to reduce tension dependent binding to F-actin reduce adherens junction localization of TRIP6 and LIMD1. The indicated cell lines expressing GFP fusions of wild-type or tension sensing mutants of TRIP6 (C) and LIMD1 (D) were stained for GFP for TRIP6 constructs or visualized using GFP fluorescence for LIMD1 constructs and either Vinculin or TRIP6 antibody to mark adherens junctions as indicated. Merged images shows TRIP6/LIMD1(green), Vinculin/TRIP6 (red), and DNA (blue). Scale bars = 50 microns.

Similarly, another study (18) observed that LIM domains function additively to increase binding to actin under strain. We tested the importance of these residues for TRIP6 and LIMD1 localization to adherens junctions. We mutated the conserved phenylalanine/tyrosine residues (F319, F377, Y449) in TRIP6 and LIMD1 (F575, F512, Y646) to alanines, individually (TRIP6) and in combination (LIMD1). We found that while individual LIM domain mutants in TRIP6 were still capable of localizing to adherens junctions, a double mutant showed reduced localization (Supplemental Figure 1A), and the triple mutant in TRIP6 (TRIP6-mut1,2,3) and LIMD1 (LIMD1-mut1,2,3) showed much reduced localization to adherens junctions compared to the wild-type proteins (Figure 4C-D).These results show that that the ability of TRIP6 and LIMD1 to localize to adherens junctions depends on the ability of each of their 3 LIM domains to sense tension.

### The ability of TRIP6 and LIMD1 LIM domains to sense tension is important for association with, and recruitment of LATS1/2 to adherens junctions

Our previous study showed that TRIP6 associated with LATS1/2 in a tension dependent manner (4). Therefore we tested whether the ability of the TRIP6 LIM domains to bind to strained actin is important for association with LATS1/2 and other tension dependent proteins at adherens junctions using a proximity labeling approach. Wild-type TRIP6, TRIP6-Δ252-277 (vinculin binding defective), and TRIP6-mut1,2,3 were fused to the promiscuous biotin ligase TurboID (20) and expressed in TRIP6-KO MCF10A cells. The TurboID protein will label proteins in close proximity with biotin. Expression of the three TurboID fusions is very similar to (TRIP6-Δ252-277 and TRIP6-mut1,2,3) or less than (wild-type TRIP6) that of endogenous TRIP6 (Supplemental Figure 1B). Staining for TRIP6 or for total biotin showed that each TurboID fusion localized similarly to its GFP fusion counterpart, with both wild-type TRIP6 and TRIP6-Δ252-277 TurboID fusions localizing to adherens junctions, and TRIP6-mut1,2,3 showing minimal staining to adherens junctions (Supplemental Figure 1C). To assess TRIP6 association with LATS1 and other adherens junctions proteins, biotinylated proteins from each cell line were purified and analyzed by western blotting. Wild-type TRIP6 and TRIP6-Δ252-277 behaved very similarly in that they both labelled LATS1, vinculin, and LIMD1 to a similar extent (Figure 5A). This labeling was specific since GAPDH (a control cytoplasmic protein) was not labeled and the adherens junction protein *α*-catenin was only weakly labeled, showing that only adherens junctions proteins in very close proximity were labeled. In contrast, TRIP6-mut1,2,3 showed almost no labelling of LATS1 and reduced labeling of vinculin and LIMD1 compared to wild-type TRIP6 and TRIP6-Δ252-277 (Figure 5A). Thus, the ability of the TRIP6 LIM domains to bind strained actin is critical for its ability to both localize to adherens junctions and bind to LATS1 and other adherens junction proteins in response to mechanical tension.

**Figure 5.**
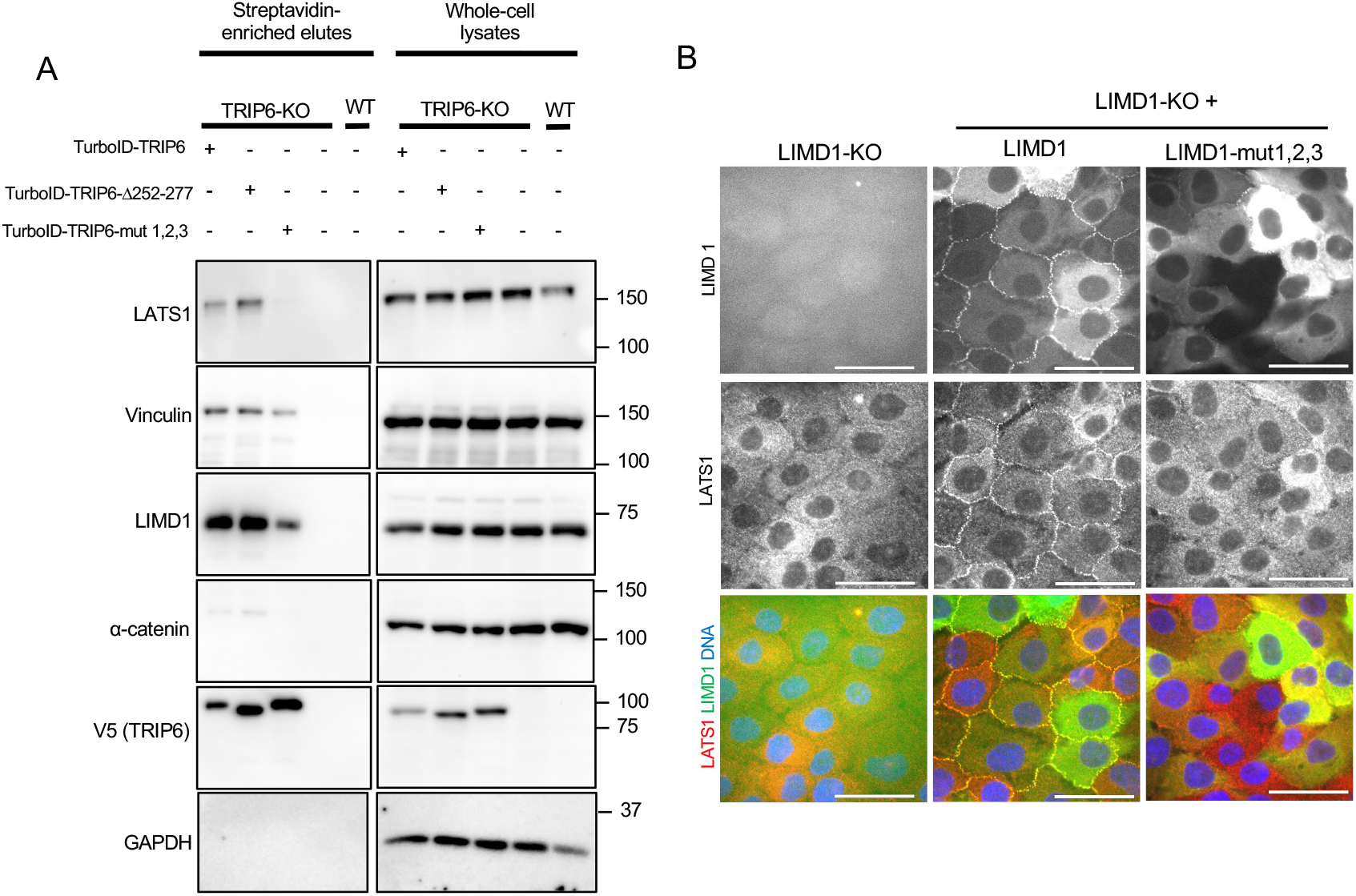
Tension dependent association of TRIP6 with LATS1 and LATS1 recruitment to adherens junctions by LIMD1 depends on the ability each protein’s LIM domains to bind f-actin under strain. (A) Protein lysates were prepared from TRIP6 knockout (TRIP6-KO) MCF10A cells expressing TurboID fusions to wild type TRIP6 or indicated TRIP6 constructs along with control wild-type MCF10A cells (WT). Biotinylated protein fractions were pulled down using streptavidin-agarose beads. Protein complexes on beads were analyzed by Western blotting using antibodies against the indicated proteins. Levels of total proteins were analyzed by blotting whole-cell lysates. (B) The indicated cell lines expressing GFP fusions of wild-type or the tension sensing mutant of LIMD1 (LIMD1-mut1,2,3) were imaged for LATS1 immunostaining signal, GFP fluorescence and DNA. Merged images show LIMD1 (green), LATS1 (red), and DNA (blue). Scale bar=50microns.

To test whether the ability to bind strained actin is also important for TRIP6 and LIMD1 recruitment of LATS1 to adherens junctions, we examined the phenotype of TRIP6 and LIMD1 knockout MCF10A cells rescued by their respective wild-type or tension sensing mutant. To our surprise, TRIP6 knockout cells (TRIP6-KO) did not show the clear defects in localization of LATS1 and LIMD1 to adherens junctions (not shown) that were previously reported using siRNA or sgRNA to knockdown TRIP6 (4, 21). The reason for this difference is not clear and was observed in multiple independent knockout clones. We suspect that the extended culturing of the cells required to isolate knockout colonies may have selected for cells that no longer need TRIP6 for tension dependent regulation of LIMD1 and LATS1. Examination of the LIMD1-KO cells showed that LATS1 localization to adherens junctions was lost in LIMD1-KO cells similar to previous observations when LIMD1 was knocked down using siRNA (5, 21). This effect is specific, since localization of TRIP6 and vinculin was not affected as observed previously (21) (Figure 5B). The defect in LATS1 localization could be rescued by expression of GFP-LIMD1 in these cells (Figure 5B). In contrast, the GFP-LIMD1-mut1,2,3 tension sensing mutant was not able to restore LATS1 localization to adherens junctions in the LIMD1-KO MCF10A cells (Figure 5B) despite being expressed at similar levels (Figure S1D). These results show that the ability of LIMD1 to bind to strained actin at adherens junctions is critical for its ability to both localize to adherens junctions and recruit LATS1 in response to mechanical tension.

## Discussion

Previous studies showed that TRIP6 and LIMD1 play a central role in tension dependent regulation of Hippo signaling (4, 5, 21). Both proteins localize to adherens junctions in a tension dependent manner, where they are required to recruit and inhibit LATS1. TRIP6 and LIMD1 can bind LATS1/2 (4, 22, 23), and TRIP6 binding has been shown to inhibit LATS1/2 by competing for binding to its activator MOB1 (4). How TRIP6 and LIMD1 are regulated by mechanical tension has remained uncertain. It was not known if these proteins sensed tension directly or were regulated by known upstream tension sensing proteins *α*-catenin and vinculin. Several lines of evidence suggested regulation by *α*-catenin and vinculin. The *α*-catenin protein is known to directly sense tension at adherens junctions, which triggers it to bind and activate vinculin and other proteins (7). Activated vinculin can then bind f-actin and other effectors (17). We showed that vinculin can bind to TRIP6 and is required for TRIP6 localization to adherens junctions and binding to LATS1/2 (4), suggesting a model where *α*-catenin is activated by tension causing it to activate vinculin, which in turn binds and activates TRIP6 causing it to localize to adherens junctions and inhibit LATS1/2. Along similar lines LIMD1 was shown to require *α*-catenin for localization to adherens junctions (5). Together these studies suggested that *α*-catenin and vinculin could be the primary sensors of mechanical strain at adherens junctions and TRIP6 and LIMD1 might not be sensors on their own but instead act downstream of *α*-catenin. However, work described here argues against this model. In the case of TRIP6, mutants that disrupt vinculin binding to TRIP6 do not have an effect on TRIP6 binding to LATS1/2 or localization to adherens junctions (Figure 2E-G). Although *α*-catenin and vinculin (in the case of TRIP6 (4)) are required for TRIP6 and LIMD1 localization to adherens junctions (Figure 1B)(5), the requirement may be through indirect mechanisms such as general effects on adherens junction assembly, regulation of the actin cytoskeleton and generation of tension. Our observation that TRIP6 binding to vinculin is not required for its recruitment to adherens junctions raises the question of what is the purpose of this binding interaction. One possibility is that the interaction could contribute to the stability/dynamics of either protein at adherens junctions. Another possibility is that TRIP6 binding could regulate vinculin. Although we did not observe an effect of TRIP6 knockdown on vinculin localization (4), another study showed that TRIP6 has a role in vinculin localization to adherens junctions (21). Further study will be required to test these possibilities.

This work showed that that the 3 LIM domains of TRIP6 and LIMD1 are both necessary and sufficient for tension dependent localization to adherens junctions. Mutations in conserved LIM domain residues, which were shown in related proteins to be required for binding to f-actin under strain (19), largely disrupted localization of TRIP6 and LIMD1 to adherens junctions (Figure 4C-D) and their ability to associate with LATS1 (in the case of TRIP6 (Figure 5A)) and recruit LATS1 (in the case of LIMD1 (Figure 5B)) to adherens junctions. Consistent with a role for binding to strained f-actin in TRIP6 and LIMD1 localization, both full length TRIP6 and LIMD1 and their respective LIM domains alone colocalized with f-actin filaments at adherens junctions (Figure 4A-B). It is curious that the LIM domains of both TRIP6 and LIMD1 localize specifically to regions at the ends of actin fibers in contact with the adherens junctions and not along the entire actin fiber. This is similar to a previous study (19) which reported that many mechano-responsive LIM domain proteins localize to focal adhesions in a manner that depends on their ability to sense tension. Together these studies suggest that regions of actin fibers close to attachment sites at focal adhesions and adherens junctions may be under higher strain than other regions along the filament.

How tension promotes association of TRIP6 and LIMD1 with LATS1/2 remains uncertain. Previous studies have shown that TRIP6, LIMD1, and related LIM domain proteins bind to LATS1/2 through their LIM domains (4, 24). For both TRIP6 and LIMD1, multiple LIM domains contribute to binding to LATS1/2, with both the first 2 LIM domains of LIMD1 being required (24), and all 3 LIM domains of TRIP6 contributing (Figure 1D). The individual LIM domains of both proteins are connected by somewhat flexible linkers, thus a binding interface involving multiple LIM domains may require specific relative positioning of the LIM domains. Therefore it is possible that binding to a strained f-actin filament may serve to align the LIM domains with respect to each other so that they can optimally bind LATS1/2. This mechanism would allow TRIP6 and LIMD1 to sense tension by binding to strained f-actin, which would align their LIM domains in an orientation favorable for LATS1/2 binding and inhibition. Future structural biology and biophysical analysis will be needed to test this model.

In summary, we found that the TRIP6 and LIMD1 proteins localize to adherens junctions through binding of their 3 LIM domains to strained f-actin at sites of filament attachment to adherens junctions. Binding to these sites is critical for tension dependent binding and recruitment of LATS1/2 to regulate Hippo pathway signaling in response to mechanical tension.

## MATERIALS AND METHODS

### Antibodies

Antibodies used for immunostaining and western blotting included rabbit anti-GFP (1:500, Cell signaling,2956), mouse anti-Vinculin (1:1000, Sigma, V-9131), mouse anti-TRIP6 (1:1000, Santa Cruz, sc-365122), rabbit anti -LATS1(1:500, Cell Signaling,3477), rabbit anti - LIMD1 (1:500, Novus biologicals, NBP2-56448), rabbit anti *α*-E-Catenin (1:500, Cell Signaling, 3236). Alexa Fluor 568 conjugated Phalloidin was used to stain for F-actin (1:100, Invitrogen, A12380). The following antibodies were used only for western blotting: mouse anti-FLAG (1:1000,Sigma,F3165), rabbit anti-GAPDH (1:1000, Cell signaling, 5174). For detection of biotinylated proteins by western blotting IRDye® 680RD Streptavidin (LI-COR Biosciences, 926-68079) was used at 1:1000.

### Immunofluorescence

MCF10A cells were cultured on coverslips and fixed with 4% Paraformaldehyde in UB (UB; 150mM NaCl, 50mM Tris pH 7.4) for 10 min, permeabilized in 0.5% Triton X-100 in UB for 5 min at room temperature, then washed 3 times in UB. They were then blocked with 10%BSA in UB for 1 hour at 20-22 °C and then incubated overnight at 4 °C with the appropriate primary antibodies. They are then washed 3 times and incubated with Alexa Flour-conjugated secondary antibodies for 1 hour at 20-22 °C. Followed by 3 more washed in UB, the coverslips were mounted onto glass slides using Prolong Gold Antifade reagent with DAPI (Invitrogen) and left to cure at room temperature overnight. The next day the slides were viewed using fluorescent microscopy and imaged. Image processing and analysis were carried out with ImageJ software.

### Cell transfection, coimmunoprecipitation and western blotting

For the coimmunoprecipitation experiments, wild type HEK293 or HEK293A (for figure 1A) cells were transfected with the indicated plasmids with Lipofectamine2000 Reagent (Invitrogen) as per manufacturer protocol. Typically, 48 hours after transfection cell lysates were prepared by adding lysis buffer (10% Glycerol (Invitrogen), 20mM Tris-HCL, pH=7, 137mM NaCl, 2mM EDTA, 1% NP 40(Invitrogen) that was supplemented with phosphatase inhibitors (2mM Sodium Vanadate,50mM Sodium Fluoride) and protease inhibitors (1mM PhenylMethylSulfonylFluoride (PMSF) and mammalian Protease Inhibitor mixture (Sigma)) directly to cells. Cells were collected using a cell scraper and pipetting back and forth. Lysates were cleared by centrifugation at 15,000 X g at 4 °C for 5 minutes and divided for use as either input or immunoprecipitation samples. Immunoprecipitation experiments were carried out using magnetic beads (Dynabeads,Invitrogen) bound to respective tag specific antibodies (detailed in figure legends) and corresponding isotype control antibody as per manufacturer’s protocol.

Following immunoprecipitation, samples were processed for Western blotting and probed using the indicated antibodies (detailed in figure legends) . To quantify Western blots following co-immunoprecipitation experiments, we performed background subtraction and densitometric analyses of respective bands using FIJI (ImageJ) and normalized to loading control (GAPDH). Data are presented as mean +/-SD (standard deviation) of three independent experiments. Student’s t-test ^(*^p<=0.05, ^**^p<=0.01, ^***^p<=0.001, ^****^p<=0.0001) was performed using GraphPad Prism software .

### Proximity Labeling

Proximity labeling methods were adapted from previous studies (25-27) with a few modifications. MCF10A-TRIP6-KO cells stably expressing lentivirally introduced TurboID fusions to TRIP6, TRIP6-Δ252-277, or TRIP6-mut 1,2,3, as well as normal and TRIP6-KO MCF10A cells (controls) were grown on 10 cm plates until 70-80% confluent. Cells were washed twice with ice-cold PBS (5 mL per 10 cm plate). Then cells were harvested by gently scraping with 1 mL of RIPA buffer (pH 7.5) [50 mM Tris–HCl (pH 7.5), 150 mM NaCl, 1% NP-40, 1 mM EDTA, 1 mM EGTA, 0.1% SDS and 0.5% sodium deoxycholate] supplemented with protease inhibitors [1mM PMSF and 1 × mammalian protease inhibitor cocktail (Millipore Sigma, P8340) and phosphatase inhibitors (1mM Na_3_VO_4_ and 10mM NaF). Harvested cells were collected into a pre-cooled 1.5 mL Eppendorf tube and cell lysates were passed through a 26-gage needle syringe on ice to eliminate cell clumps. Cell lysates were then sonicated at 4 °C for three 10 second bursts with 2-minute pauses using a Diagenode Picoruptor sonicator (P-190705). Then each cell lysate was treated with 100 units of benzonase nuclease (Millipore Sigma, E1014) and incubated at 4°C with shaking for an hour. A total of 110 *μ* L from each sample was saved for BCA protein quantification and whole cell lysate analysis via SDS-PAGE. For SDS-PAGE analysis 100 *μ* L of sample was mixed with 1 × Laemmli SDS-sample buffer and boiled at 95°C for 5 minutes. The remaining cell lysates were cleared by centrifugation at full speed in a microcentrifuge at 4 °C for 10 minutes. The resultant superannuants were incubated with pre-washed streptavidin-agarose beads (Pierce™ High-Capacity Streptavidin Agarose, Thermo Fisher, 20359) derived from 100 *μ* L of bead slurry and incubated overnight at 4 °C with shaking (the beads were washed three times with 1 mL of RIPA buffer by pelleting the beads at 400 g for 1 minute in a microcentrifuge and aspirating off the supernatant before adding the next wash). Following the overnight incubation, the beads were pelleted at 400 g for 1 minute at 4 °C in a microcentrifuge. To remove unbound proteins the beads were washed sequentially with RIPA buffer (pH 7.5) (2×), with 1M KCl (1×), with 50 mM (NH_4_) HCO_3_ (pH 8.0) (3x), with × RIPA buffer (pH 7.5) (2×), and with × 50 mM Tris HCl (pH 7.4) (3x). All washes were done at 4 °C for 3 minutes on a shaker with 1ml buffer supplemented with protease and phosphatase inhibitors and 400 g spins for one minute. Proteins bound to the beads were removed by adding 1 × Laemmli SDS-sample buffer saturated with 10 mM biotin and boiling at 95°C for 5 minutes.

Both the whole cell lysate and eluted protein fractions were separated on 10% (w/v) SDS PAGE resolving gel followed by Western blot analysis for various proteins via ECL method or for biotinylated proteins via Li-Cor Odyssey Infrared Imaging method.

### Cell Culture

Human embryonic kidney (HEK293, HEK293A) cell lines were grown in Dulbeco’s Modified Eagle medium (DMEM, Gibco) supplemented with 10% (v/v) fetal bovine serum (FBS, Gibco) and 1% (v/v) Penicillin/Streptomycin (Invitrogen). The MCF10A human mammary epithelial cell line was cultured in DMEM/F12 (1:1) media supplemented with 5% (v/v) horse serum (Gibco), 20 ng/ml epidermal growth factor (Peprotech), 0.5 *μ*g/ml Hydrocortisone (Sigma), 100ng/ml Cholera toxin (Sigma), 10*μ*g/ml Insulin (Sigma) and 1% (v/v) Penicillin/Streptomycin (Invitrogen). All cell lines were cultured in a humidified incubator that is maintained at 37 °C with 5% CO2. TRIP6 or LIMD1 knockout (TRIP6-KO/LIMD1-KO) MCF10A cell lines that are stably expressing the indicated constructs were generated by lentivirus infection followed by selection with Puromycin (Gibco, at working concentration 1*μ*g/ml). Virus packaging and supernatant preparation were carried out by RNAi Core Facility (UMASS Chan).

### CRISPR mediated TRIP6 and LIMD1 knock out (KO) cell line development

MCF10A cells stably expressing Cas9 were transfected with TRIP6 or LIMD1 specific multi-guide sgRNA pools that were purchased from Synthego. Transfections were carried out in a 12 well dish using Invitrogen Lipofectamine RNAiMax kit as per manufacturer’s protocol. Three days following transfection cells were processed for western blot to test for overall knock down efficiency and also heavily diluted with conditioned media and plated on collagen coated dishes to derive clones of for each knockout. Individual colonies were picked using sterile pyrex rings one end of which was dipped into sterile silicone grease and pressed to bottom of the culture dish to create an isolated well of cells. Isolated colonies were then trypsin treated to dislodge cells and propagated on collagen coated culture vessels at optimal confluency using conditioned media to facilitate cell attachment and growth. Colonies were screened for TRIP6 and LIMD1 expression using western blot and immunostaining. Colonies that showed no bands by western blot and no staining by immunofluorescence were saved. In the event that the first round of colony selection resulted in mixed clones cells with and without TRIP6 or LIMD1 expression, they were processed for additional rounds of clone picking and subsequent screening as described above to obtain clonal knock out lines.

---

**Supplementary Figure 1.**
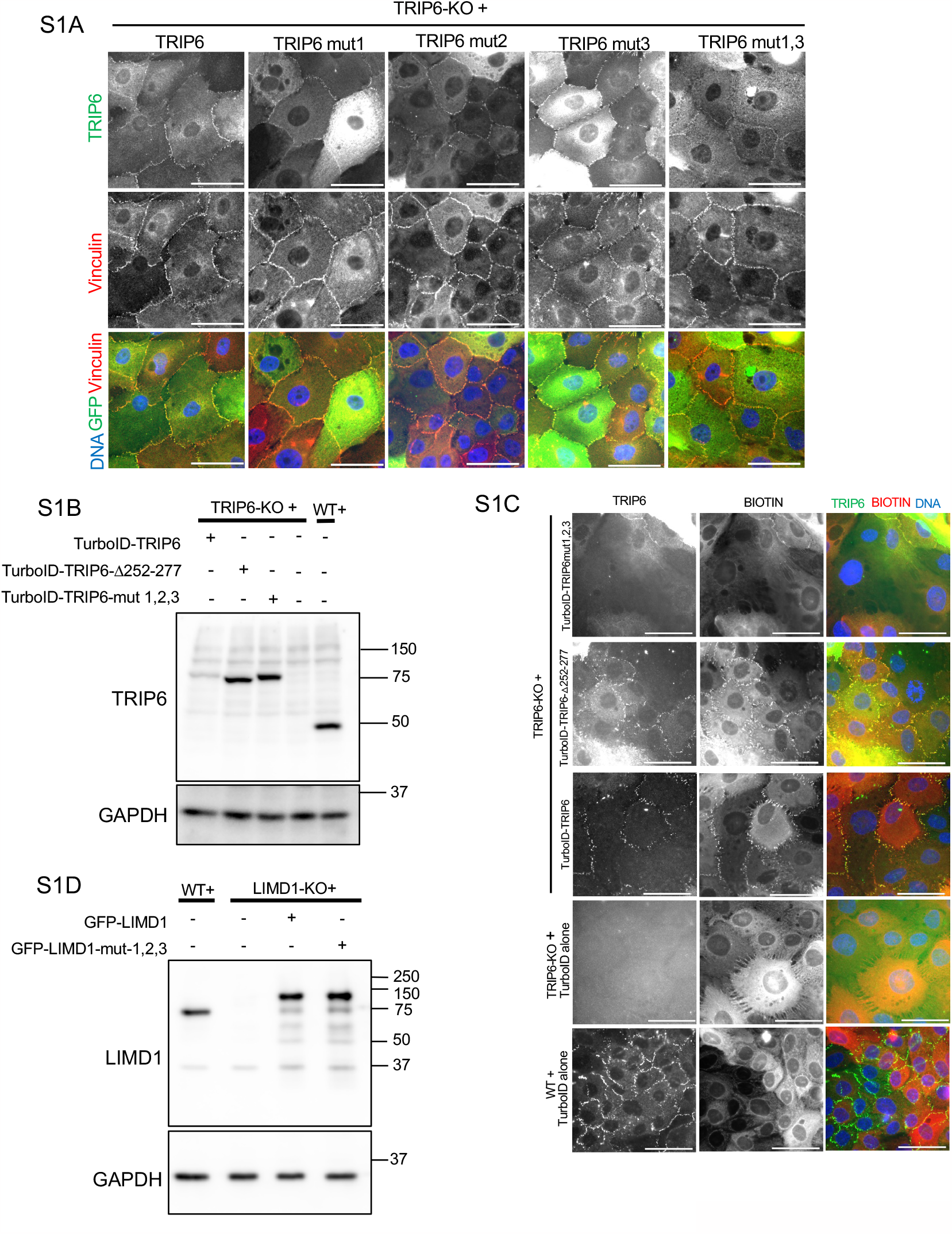
(A) TRIP6 knockout (TRIP6-KO) MCF10A cells stably expressing (via lentiviral transduction) GFP fusions of wild-type TRIP6 (TRIP6) or the indicated tension sensing mutants in each LIM domain were stained for GFP and Vinculin. Merged images show TRIP6 (green), Vinculin (red), and DNA (blue). Scale bars=50microns. (B) Expression of TurboID fusions to indicated TRIP6 constructs was analyzed by western blotting from whole cell lysates of wild-type MCF10A cells (WT) and TRIP6 knockout (TRIP6-KO) MCF10A cells stably expressing the indicated constructs of TRIP6. Blots were probed using antibodies against TRIP6 and GAPDH. (C) Localization of TurboID fusions to the indicated TRIP6 constructs was analyzed by immunostaining of TRIP6 knockout (TRIP6-KO) or wild-type MCF10A cells expressing indicated TurboID tagged TRIP6 constructs or TurboID alone as indicated. Cell lines were stained with antibodies against TRIP6 and fluorescent streptavidin to label biotin (BIOTIN). Merged images shows TRIP6 (green), Biotin (red), and DNA (blue). Scale bars=50microns. (D) Expression levels of LIMD1 in wild type MCF10A cells or LIMD1-KO MCF10A cells stably expressing wild-type (GFP-LIMD1) or the tension sensing mutant (GFP-LIMD1-mut 1,2,3) of LIMD1. Whole cell lysates were probed by Western blotting using antibodies against LIMD1 and GAPDH.

## REFERENCES

1. Mammoto T, Mammoto A, Ingber DE. Mechanobiology and developmental control. Annu Rev Cell Dev Biol. 2013;29:27–61. Epub 2013/10/09. doi: 10.1146/annurev-cellbio-101512-122340. PubMed PMID: 24099083.

2. DuFort CC, Paszek MJ, Weaver VM. Balancing forces: architectural control of mechanotransduction. Nat Rev Mol Cell Biol. 2011;12(5):308–19. Epub 2011/04/22. doi: 10.1038/nrm3112. PubMed PMID: 21508987; PMCID: PMC3564968.

3. Rauskolb C, Sun S, Sun G, Pan Y, Irvine KD. Cytoskeletal tension inhibits Hippo signaling through an Ajuba-Warts complex. Cell. 2014;158(1):143–56. Epub 2014/07/06. doi: 10.1016/j.cell.2014.05.035S0092-8674(14)00722-3 [pii]. PubMed PMID: 24995985; PMCID: PMC4082802.

4. Dutta S, Mana-Capelli S, Paramasivam M, Dasgupta I, Cirka H, Billiar K, McCollum D. TRIP6 inhibits Hippo signaling in response to tension at adherens junctions. EMBO Rep. 2018;19(2):337–50. Epub 2017/12/10. doi: 10.15252/embr.201744777. PubMed PMID: 29222344; PMCID: PMC5797958.

5. Ibar C, Kirichenko E, Keepers B, Enners E, Fleisch K, Irvine KD. Tension-dependent regulation of mammalian Hippo signaling through LIMD1. J Cell Sci. 2018;131(5). Epub 2018/02/15. doi: 10.1242/jcs.214700. PubMed PMID: 29440237; PMCID: PMC5897721.

6. Dasgupta I, McCollum D. Control of cellular responses to mechanical cues through YAP/TAZ regulation. J Biol Chem. 2019. Epub 2019/10/09. doi: 10.1074/jbc.REV119.007963. PubMed PMID: 31594864.

7. Pokutta S, Weis WI. Structure and mechanism of cadherins and catenins in cell-cell contacts. Annu Rev Cell Dev Biol. 2007;23:237–61. Epub 2007/06/02. doi: 10.1146/annurev.cellbio.22.010305.104241. PubMed PMID: 17539752.

8. Yonemura S, Wada Y, Watanabe T, Nagafuchi A, Shibata M. alpha-Catenin as a tension transducer that induces adherens junction development. Nat Cell Biol. 2010;12(6):533–42. Epub 2010/05/11. doi: 10.1038/ncb2055ncb2055 [pii]. PubMed PMID: 20453849.

9. Choi HJ, Pokutta S, Cadwell GW, Bobkov AA, Bankston LA, Liddington RC, Weis WI. alphaE-catenin is an autoinhibited molecule that coactivates vinculin. Proc Natl Acad Sci U S A. 2012;109(22):8576–81. doi: 10.1073/pnas.1203906109. PubMed PMID: 22586082; PMCID: PMC3365184.

10. Kim TJ, Zheng S, Sun J, Muhamed I, Wu J, Lei L, Kong X, Leckband DE, Wang Y. Dynamic visualization of alpha-catenin reveals rapid, reversible conformation switching between tension states. Curr Biol. 2015;25(2):218–24. Epub 2014/12/30. doi: 10.1016/j.cub.2014.11.017S0960-9822(14)01444-4 [pii]. PubMed PMID: 25544608; PMCID: PMC4302114.

11. Yao M, Qiu W, Liu R, Efremov AK, Cong P, Seddiki R, Payre M, Lim CT, Ladoux B, Mege RM, Yan J. Force-dependent conformational switch of alpha-catenin controls vinculin binding. Nat Commun. 2014;5:4525. Epub 2014/08/01. doi: 10.1038/ncomms5525. PubMed PMID: 25077739.

12. Wen KK, Rubenstein PA, DeMali KA. Vinculin nucleates actin polymerization and modifies actin filament structure. J Biol Chem. 2009;284(44):30463–73. Epub 2009/09/09. doi: 10.1074/jbc.M109.021295. PubMed PMID: 19736312; PMCID: PMC2781601.

13. le Duc Q, Shi Q, Blonk I, Sonnenberg A, Wang N, Leckband D, de Rooij J. Vinculin potentiates E-cadherin mechanosensing and is recruited to actin-anchored sites within adherens junctions in a myosin II-dependent manner. J Cell Biol. 2010;189(7):1107–15. Epub 2010/06/30. doi: 10.1083/jcb.201001149. PubMed PMID: 20584916; PMCID: PMC2894457.

14. Cohen DM, Chen H, Johnson RP, Choudhury B, Craig SW. Two distinct head-tail interfaces cooperate to suppress activation of vinculin by talin. J Biol Chem. 2005;280(17):17109–17. doi: 10.1074/jbc.M414704200. PubMed PMID: 15728584.

15. Vite A, Zhang C, Yi R, Emms S, Radice GL. alpha-Catenin-dependent cytoskeletal tension controls Yap activity in the heart. Development. 2018;145(5). Epub 2018/02/23. doi: 10.1242/dev.149823. PubMed PMID: 29467248; PMCID: PMC5868989.

16. Hirota T, Morisaki T, Nishiyama Y, Marumoto T, Tada K, Hara T, Masuko N, Inagaki M, Hatakeyama K, Saya H. Zyxin, a regulator of actin filament assembly, targets the mitotic apparatus by interacting with h-warts/LATS1 tumor suppressor. J Cell Biol. 2000;149(5):1073–86. Epub 2000/06/01. PubMed PMID: 10831611; PMCID: PMC2174824.

17. Bays JL, DeMali KA. Vinculin in cell-cell and cell-matrix adhesions. Cell Mol Life Sci. 2017. doi: 10.1007/s00018-017-2511-3. PubMed PMID: 28401269.

18. Winkelman JD, Anderson CA, Suarez C, Kovar DR, Gardel ML. Evolutionarily diverse LIM domain-containing proteins bind stressed actin filaments through a conserved mechanism. Proc Natl Acad Sci U S A. 2020;117(41):25532–42. Epub 2020/09/30. doi: 10.1073/pnas.2004656117. PubMed PMID: 32989126; PMCID: PMC7568268.

19. Sun X, Phua DYZ, Axiotakis L, Jr., Smith MA, Blankman E, Gong R, Cail RC, Espinosa de Los Reyes S, Beckerle MC, Waterman CM, Alushin GM. Mechanosensing through Direct Binding of Tensed F-Actin by LIM Domains. Dev Cell. 2020;55(4):468–82 e7. Epub 2020/10/16. doi: 10.1016/j.devcel.2020.09.022. PubMed PMID: 33058779; PMCID: PMC7686152.

20. Branon TC, Bosch JA, Sanchez AD, Udeshi ND, Svinkina T, Carr SA, Feldman JL, Perrimon N, Ting AY. Efficient proximity labeling in living cells and organisms with TurboID. Nat Biotechnol. 2018;36(9):880–7. Epub 2018/08/21. doi: 10.1038/nbt.4201. PubMed PMID: 30125270; PMCID: PMC6126969.

21. Venkatramanan S, Ibar C, Irvine KD. TRIP6 is required for tension at adherens junctions. J Cell Sci. 2021;134(6). Epub 2021/02/10. doi: 10.1242/jcs.247866. PubMed PMID: 33558314; PMCID: PMC7970510.

22. Das Thakur M, Feng Y, Jagannathan R, Seppa MJ, Skeath JB, Longmore GD. Ajuba LIM proteins are negative regulators of the Hippo signaling pathway. Curr Biol. 2010;20(7):657–62. Epub 2010/03/23. doi: S0960-9822(10)00223-X [pii] 10.1016/j.cub.2010.02.035. PubMed PMID: 20303269; PMCID: PMC2855397.

23. Sun G, Irvine KD. Ajuba family proteins link JNK to Hippo signaling. Sci Signal. 2013;6(292):ra81. Epub 2013/09/12. doi: 10.1126/scisignal.20043246/292/ra81 [pii]. PubMed PMID: 24023255; PMCID: PMC3830546.

24. Jagannathan R, Schimizzi GV, Zhang K, Loza AJ, Yabuta N, Nojima H, Longmore GD. AJUBA LIM Proteins Limit Hippo Activity in Proliferating Cells by Sequestering the Hippo Core Kinase Complex in the Cytosol. Mol Cell Biol. 2016;36(20):2526–42. Epub 2016/07/28. doi: 10.1128/MCB.00136-16. PubMed PMID: 27457617; PMCID: PMC5038147.

25. Cho KF, Branon TC, Udeshi ND, Myers SA, Carr SA, Ting AY. Proximity labeling in mammalian cells with TurboID and split-TurboID. Nat Protoc. 2020;15(12):3971–99. Epub 2020/11/04. doi: 10.1038/s41596-020-0399-0. PubMed PMID: 33139955.

26. Roux KJ, Kim DI, Burke B. BioID: a screen for protein-protein interactions. Curr Protoc Protein Sci. 2013;74:19 23 1-19 23 14. Epub 2014/02/11. doi: 10.1002/0471140864.ps1923s74. PubMed PMID: 24510646.

27. Lambert JP, Tucholska M, Go C, Knight JD, Gingras AC. Proximity biotinylation and affinity purification are complementary approaches for the interactome mapping of chromatin-associated protein complexes. J Proteomics. 2015;118:81–94. Epub 2014/10/05. doi: 10.1016/j.jprot.2014.09.011. PubMed PMID: 25281560; PMCID: PMC4383713.

